# Comparative Single-Cell Transcriptomics Uncovers Shared and Distinct Molecular Signatures in Cystic Fibrosis and Primary Ciliary Dyskinesia

**DOI:** 10.1101/2025.11.17.688876

**Authors:** Nicholas Hadas, Huihui Xu, Wang Kyaw Twan, Shambhawee Neupane, Ahmed Elgamal, Jeffrey R Koenitzer, Amjad Horani

## Abstract

**Rational:** Cystic Fibrosis (CF) and Primary Ciliary Dyskinesia (PCD) are both inherited respiratory disorders that result in impaired mucociliary clearance, and chronic sinopulmonary disease. Although the current approach to PCD management is extrapolated from CF care, both conditions arise from distinct genetic and molecular mechanisms.

**Methods:** Here we performed a comparative transcriptomic analysis between CF and PCD to compare the cellular heterogeneity, molecular pathways and gene networks differences using publicly available sequencing data as well as those performed by our group. To explore gene regulatory networks, a pre-trained transformer model (scGPT) was fine-tuned using an integrated dataset, and differential attention analysis was conducted to identify genes and pathways with altered attention scores between the two conditions.

**Results:** The comparative transcriptomic analysis revealed distinct molecular signatures between PCD and CF, which differed from normal cells. In ciliated cells, differential gene expression and pathway investigation highlighted the NRF2 pathway’s considerable overrepresentation in PCD compared to CF and healthy conditions. This observation was further supported by scGPT analysis, which revealed increased incoming attention to the NRF2 pathway markers. In secretory cells, PCD and CF exhibited increased immune and inflammatory signaling compared to controls. While similar inflammatory processes were active, results suggested a stronger inflammatory pattern in CF secretory cells compared to PCD and confirmed the activation of the unfolded protein response (UPR) pathway.

**Conclusion:** These findings highlight the different molecular signatures between both conditions and the need for unique approaches to management in PCD compared to CF.

## Introduction

Cystic Fibrosis (CF) and Primary Ciliary Dyskinesia (PCD) are rare, inherited disorders that are primarily associated with respiratory complications attributed to impaired mucociliary clearance (MCC) (1). Current approaches to PCD care stems from experience managing individuals with CF, as well as other suppurative lung diseases and idiopathic bronchitis. Recent insight into the molecular mechanisms of both conditions highlights differences between CF and PCD (1). In CF, a monogenic disorder caused by pathogenic variants in the cystic fibrosis transmembrane conductance regular (*CFTR*) gene, mucociliary disruption is caused by an altered mucus layer composition overlaying motile cilia (2, 3). PCD in comparison is caused by variants in over 60 cilia-associated genes, yielding dysfunctional or absent cilia motility and less severely affected mucus layer (4).

Recent advances in single cell RNA sequencing (scRNA-seq) have increased our ability to query the transcriptional profile of cells in different disease conditions, revealing new insights into perturbed pathways including in PCD and CF (5–7). These valuable findings are crucial for understanding disease mechanisms and potential new avenues for targeted treatments (6–8). While there are several studies that have individually investigated both diseases with transcriptomic analysis tools, comparative transcriptomic analyses between CF and PCD have not been extensively investigated. This is important as differences in pathway activation may yield insight onto potential targeted therapy, that may be effective in one but not the other condition.

We recently reported increased expression of inflammatory pathways in multiciliated cells retrieved from individuals with PCD (7, 9). We found that multiciliated cells from variants in the address proteins *CCDC39* and *CCDC40*, as well as the motor dynein protein *DNAH5* result in differential gene expression of pathways associated with proteostasis, cell stress, and oxidative phosphorylation. Increased cellular stress was not limited to multiciliated cells but was also identified in secretory cells suggesting possible cross talk between different cell types, or a disease state affecting non-ciliated cells through a generalized mucociliary defect. Moreover, our recent findings suggest that multiciliated PCD cells activate protective mechanisms including targets of the NRF2 pathway (7).

To further elucidate the similarities and differences in cellular heterogeneity and cellular pathways between CF and PCD, we compared the expression profiles between both conditions using single cell RNA sequencing (scRNA-seq). Nasal cells from PCD and CF patients were cultured *in vitro* before single cell analysis (7, 9) and compared to data from two CF published datasets that also incorporated air-liquid interface growth protocols (8, 10).

Our findings suggest activation of similar pathways in these two conditions, with unique pathways that are activated in PCD and CF likely due to different molecular mechanisms dominant in each condition. These findings further emphasize the need for unique management approaches in PCD compared to CF.

## Results

### Airway epithelial cell differentiation is influenced by the in vitro culture system

Nasal epithelial cells were collected from healthy individuals and those with PCD or CF. Five healthy individuals and seven patients with confirmed PCD with pathogenic variants in *DNAH5, CCDC39* or *CCDC40* were included in this analysis, as previously reported (7, 9). Additionally, nasal epithelial cells were collected from two individuals with homozygous *CFTR* c.1521_1523del (p.F508del) causative of cystic fibrosis (**Supplemental Table 1**). All samples were grown using air-liquid interface (ALI) conditions and sequenced as we described previously (11, 12). For comparison, we also retrieved publicly available data from two CF series that included airway cells from 5 CF patients and 5 unrelated healthy individuals grown in ALI conditions (7, 9) (**Figure 1A**).

**Figure 1.**
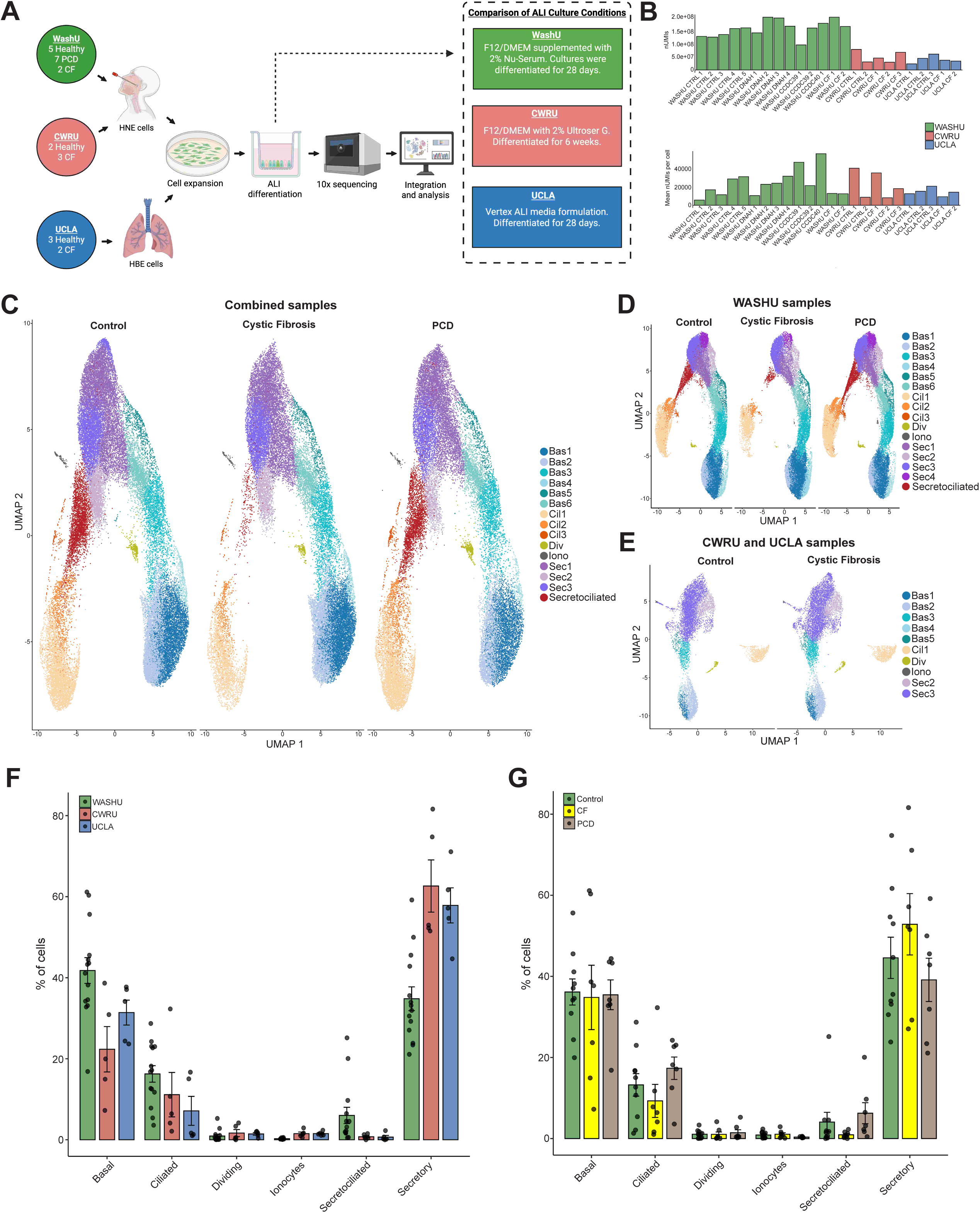
Single-cell dataset comparison and primary cell clustering. **(A)** Schematic overview of experimental design **(B)** Quantification of total and mean unique molecular identifiers (UMIs) per individual sample. **(C)** Uniform Manifold Approximation and Projection (UMAP) visualization of integrated airway cells from all three cohorts. Cells are split by condition (Control vs CF vs PCD) and colored by annotated subcluster, identifying basal, secretory, secretociliated, dividing and ionocyte populations. **(D)** UMAP visualization of the integrated CWRU and UCLA public dataset, stratified by condition (Control vs. CF). **(E)** UMAP visualization of the WashU dataset, stratified by condition (Control vs. PCD vs. CF). **(F)** Bar plot quantifying the proportional distribution of major cell types across institutions in the integrated dataset. Error bars represent the standard error of the mean (SEM). n for WASHU = 14 biologically independent samples, n = 5 for CWRU, n = 5 for UCLA. **(G)** Bar plot quantifying the proportion of major cell types, stratified by disease condition. Error bars represent SEM. Where, n = 10 biologically independent samples for control, n = 7 for CF and n = 7 for PCD.

To approximate and investigate differences in sequencing depth, the total number of unique molecular identifiers (UMIs) and mean UMIs per cell were visualized across samples. Samples collected at the Washington University site, on average had higher numbers of UMIs per cells, suggesting higher sequencing depth per cell (**Figure 1B**). Unsupervised clustering and UMAP analysis of the combined scRNA-seq data identified expected airway epithelial populations, including basal (Bas), secretory (Sec), multiciliated (Cil), secretociliated, ionocytes (Iono) and dividing cells (Div) (**Figure 1C**) and was guided by the expression of canonical markers (**Supplemental Figure 1A**) (13, 14). Multiciliated cells were further subdivided into subclusters that likely represent different stages of differentiation, as we previously reported, including late mature ciliated cells (*FOXJ1*+, *DNAH7*+, high *CFAP54*), early mature ciliated cells (*FOXJ1*+, low *CFAP54*), and deutorostomal cells (*FOXJ1*+, *CCNO*+, *DEUP1*+, *PLK4*+) (7). Similarly, the remaining major cell types were also resolved into subclusters, each characterized by a transcriptional signature as determined by differential gene expression analysis (**Supplemental Figure 1B)**.

In comparison to WASHU generated samples, those retrieved from UCLA and CWRU datasets had less multiciliated cells (**Figure 1D**, **Figure 1E)** and lacked a secretociliated cell cluster. Moreover, UCLA and CWRU datasets had a higher proportion of secretory cells than multiciliated cells (**Figure 1F and Supplemental Figure 2B)**. This is likely attributed to the use of different culture media to derive basal progenitor cells. This observation is consistent with a previous report showing that specific composition of culture media can influence the balance between basal cells proliferation and their differentiation into mature cell types (15). However, the cell type proportions were more similar when directly compared across the different conditions (**Figure 1G**).

A deeper comparative analysis between the local WASHU generated and public CF samples confirmed the changes in cell composition (**Supplemental Figure 2A and Supplemental Figure 2B**), revealed divergent pathways in secretory cells involving interferon signaling and antigen processing (**Supplemental Figure 2C and Supplemental Figure 2D**) and differing number of DEGs between their controls (**Supplemental Figure 2E**).

### PCD cells have unique transcriptional signature in multiciliated cells compared to CF

To evaluate the differences between cellular responses of PCD multiciliated cells and those of CF, we performed a comparative analysis of the transcriptional data. Compared to controls, multiciliated PCD cells had increased expression of pathways associated with inflammation including complement, as well as the NRF2 pathway, as we have previously shown (**Figure 2A and 2B**) (7). In prior work, we have hypothesized that PCD cells may have an increased oxidative burden necessitating activation of the NRF2 pathway and *GSTA2*, a highly regulated NRF2 target (7). Several NRF2 targets and genes related to oxidative stress protection were differentially expressed in multiciliated cells PCD when compared to CF (**Figure 2C and 2D**). Moreover, the number of cells expressing *GSTA2* and the overall expression levels of *GSTA2* were markedly elevated in PCD compared to CF and healthy control cells, particularly among multiciliated and secretociliated clusters (**Figure 2E, 2F and 2H**). These results further emphasize *GSTA2*’s expression and activity in multiciliated cells. Conversely, the protective NRF2 pathway was less active in CF cells. To assess the combined activity of NRF2-activated genes, we calculated a module score, which aggregates the expression levels of Reactome’s curated cytoprotective NRF2 genes into a single metric (**Figure 2G**) (16). The NRF2 module score was lower in CF and controls compared to PCD across ciliated, secretory and secretociliated cells, indicating decreased NRF2 activity (**Figure 2G**). This diminished NRF2 signature in CF ciliated cells was reinforced by significantly decreased levels of *GSTA2* and *ALDH3A1* in CF compared to controls (**Figure 2I**). This suggests that NRF2’s role and cellular mechanisms for managing oxidative stress differs between the two conditions. While key downstream NRF2 activated targets demonstrated elevated overall aggregated expression levels in PCD multiciliated cells, distinct PCD genotype groups displayed varying signatures, with the *DNAH5* variant displaying the greatest expression on average (**Figure 2I**). Furthermore, a pathway analysis of the upregulated DEGs in CF ciliated cells versus control established the absence of NRF2 activity (**Supplemental Figure 3A**).

**Figure 2.**
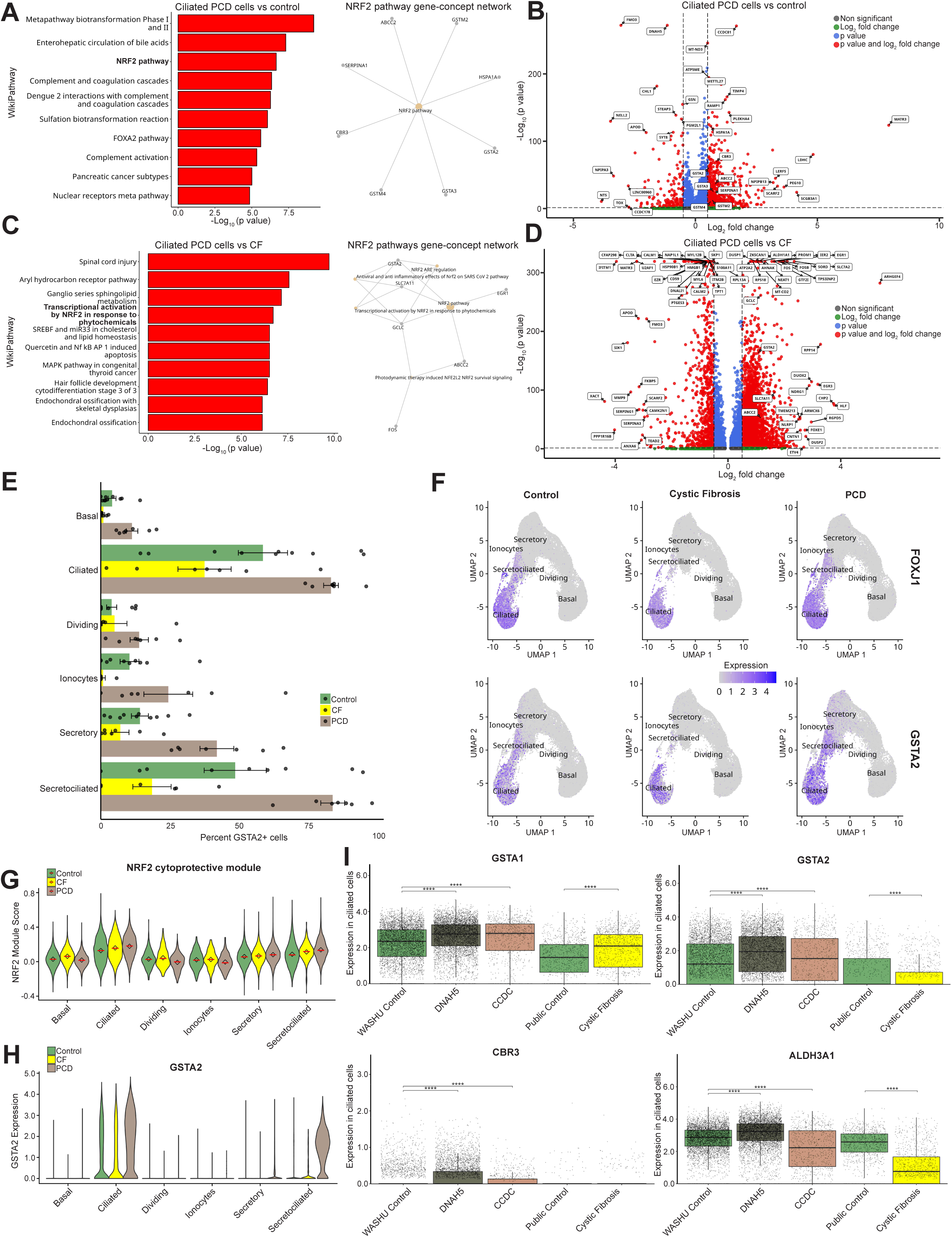
Comparative expression of GSTA2 and NRF2 markers. **(A)** Overrepresentation analysis (ORA) of WikiPathways for upregulated differentially expressed genes (DEGs) in ciliated PCD cells versus control. The top 10 enriched pathways are shown. Bottom panel depicts gene-concept network illustrating contributing DEGs for the NRF2 pathway. **(B)** Volcano plot illustrating upregulated DEGs in ciliated PCD cells versus control, with top DEGs and key NRF2-related genes highlighted. Significance cutoffs were set at an adjusted p-value (p-value < 0.05) and Log_2_ fold change > 0.5, respectively. **(C)** ORA of WikiPathways for DEGs in ciliated PCD cells versus CF. A gene-concept network display significant NRF2-related pathways (p < 0.05) in bottom panel. **(D)** Volcano plot of DEGs in ciliated PCD cells versus CF, annotated with upregulated NRF2-related genes. **(E)** Bar plot quantifying the proportion of *GSTA2*-expressing cells across conditions and cell subclusters. Error bars represent SEM. **(F)** UMAP visualizations illustrating *FOXJ1* (ciliated cell marker) and *GSTA2* expression levels, stratified by condition. **(G)** Violin plots of the ‘NRF2 cytoprotective signaling’ gene module score (WikiPathways). Red lines indicate the median; black diamonds indicate the mean. **(H)** Violin plots quantifying *GSTA2* expression across all major cell types and conditions. **(I)** Violin plots showing average expression of selected NRF2-related genes in ciliated cells, stratified by condition and PCD genotype. Significance determined by pairwise Wilcoxon rank-sum tests. Statistical significance was assessed using the pairwise Wilcoxon rank-sum test. Whiskers indicate p-value significance levels: *p < 0.05, **p < 0.01, ***p < 0.001, ****p < 0.0001; ns, not significant.

#### PCD and CF have shared and distinct immune signaling patterns

Transcriptional analysis of the ALI-differentiated epithelial cells allows investigation of intrinsic epithelial immune responses in a controlled environment, largely independent of environmental factors. We first compared the heightened immune and inflammatory state in PCD and CF versus controls through differential gene expression analysis, revealing shared groups of upregulated genes, which included chemokines and cytokines, interferons and ISGs, damage-associated molecular patterns (DAMPs) and antimicrobial markers (**Figure 3A).** Analysis of the immune-related markers revealed both overlapping and unique profiles of upregulated genes between the two diseases, with much of this activity driven by the secretory cells. Notably, prominent pro-inflammatory cytokines, including *CXCL8*/*IL-8* and *CXCL5*, were upregulated in the secretory cells of both conditions. (**Figure 3A**). The chronic inflammatory state in the secretory cells was further confirmed through pathway analysis, uncovering neutrophil degranulation as a significant shared pathway (**Figure 3B and 3C**). Additionally, bacterial peptides and chemokine receptors bind chemokines were overrepresented in PCD, while pathways related to proinflammatory eicosanoid activity and interleukin-13 signaling were identified in cystic fibrosis secretory cells (**Figure 3B and 3C**). To assess the chronic inflammation driven by neutrophil-recruiting cytokines, we calculated an aggregated expression score for a module of relevant genes from the Reactome database (16). This analysis confirmed that the cytokine signaling pathway was more active in diseased cells across major cell types, with a particularly strong signal in cystic fibrosis.

**Figure 3.**
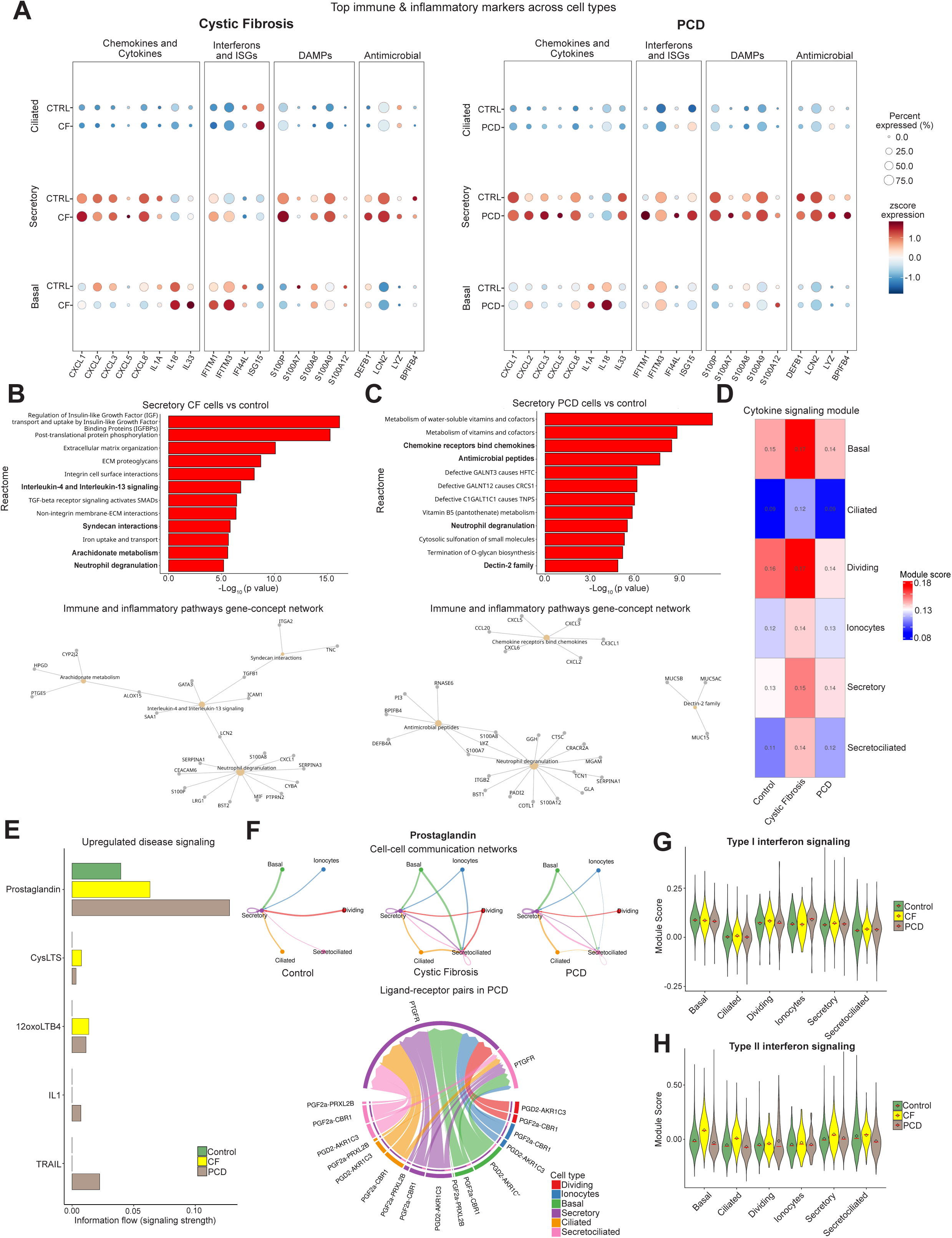
Comparative inflammatory marker expression and signaling. **(A)** Dot plot illustrating the z-scored average expression and percentage of cells expressing top inflammatory DEGs. Markers were identified by Wilcoxon rank-sum test (diseased vs. control) and filtered by Log_2_ fold change and adjusted p-value (p < 0.05). Left: Public CF versus control. Right: WashU PCD versus control. **(B)** ORA of Reactome pathways for upregulated DEGs in secretory CF cells versus control. The top 12 pathways are shown. A gene-concept network highlights contributing genes for enriched immune pathways in the bottom panel. **(C)** ORA of Reactome pathways for upregulated DEGs in secretory PCD cells versus control. Bottom panel depicts a gene-concept network highlights contributing genes for enriched immune pathways. **(D)** Heatmap displaying the average ’Cytokine Signaling in Immune System’ (Reactome) module score across all major cell types and conditions. **(E)** Comparative analysis of overall information flow (interaction strength) for upregulated immune and inflammatory signaling pathways in PCD and CF versus control, as inferred by CellChat. **(F)** CellChat analysis of prostaglandin signaling. Top: Circle plots visualizing the inferred intercellular signaling networks across conditions. Bottom: Chord diagram detailing the specific ligand-receptor pairs contributing to the prostaglandin network in PCD. **(H)** Violin plots quantifying the Type I Interferon module score (WikiPathways) across conditions. **(I)** Violin plots quantifying the Type II Interferon module score (WikiPathways) across conditions. For **(H)** and **(I)**, red lines indicate the median; black diamonds indicate the mean.

Analysis of cell-to-cell communication with CellChat, uncovered specific inflammatory communication networks with increased interaction strength in PCD and CF compared to healthy control cells (17). These selected interactions were dominated by eicosanoid signaling (**Figure 3E**). Both PCD and CF had increased signaling of prostaglandins, IL1, Leukotriene B4 and TNF-related apoptosis-inducing ligand (TRAIL) compared to healthy cells. While elevated in both disease conditions, CF cells had increased Leukotriene B4 signaling compared to PCD, further emphasizing the role of neutrophil inflammation in CF versus PCD. The dysregulation of the arachidonic metabolism in cystic fibrosis airway cells and the resulting excess levels of prostaglandins and leukotrienes may be contributing to unresolved airway inflammation (18, 19). A similar pattern may be at play in PCD. The prostaglandin signaling pathway exhibited expanded intercellular communication in CF and PCD, and was characterized by active interactions between the PGF_2_α ligand and its receptor *PTGFR* in secretory PCD cells (**Figure 3F**).

Interferon signaling was more prominent in CF versus PCD. To further elucidate drivers of the immune response in PCD and cystic fibrosis, we calculated module scores to summarize the activity of key interferon signaling pathways from the WikiPathways database (20) (**Figure 3G and 3H**). Reflecting the characteristic elevated inflammatory transcriptional state, cystic fibrosis cells exhibited strong upregulation of type II interferon signaling across all major cell types, and moderately augmented interferon type I signaling in secretory cells. In contrast, PCD did not appear to be overly driven by interferon-mediated mechanisms (**Figure 3G and 3H**). Furthermore, interferon signaling was an enriched pathway in the direct comparison between CF and PCD in both ciliated and secretory cells (**Supplemental Figure 3A and 3B**), suggesting that a heightened epithelial interferon response is a driver of the CF inflammatory profile and could be a potential differentiator from PCD. Immune pathways were not apparent when directly comparing PCD to CF secretory cells (**Supplemental 3C**).

#### Gene regulatory network analysis shows the prominent role of the NRF2 pathway in PCD

To define the inferred gene regulatory networks (GRNs) that drive each disease state using an unbiased approach, we fined-tuned and processed scGPT’s 2.1 million lung cell transformer model with our integrated dataset (**Figure 4A**) (21). The model was trained by randomly masking the expression of certain genes and learning to predict them based on the context of the remaining genes. This stimulated the transformer model to learn complex, predictive relationships across the transcriptome. The strength of these relationships was quantified by “attention scores”. Differential attention matrices were then generated to characterize gene-gene interactions that are stronger in each disease state compared to controls, on a per cell-type basis.

**Figure 4.**
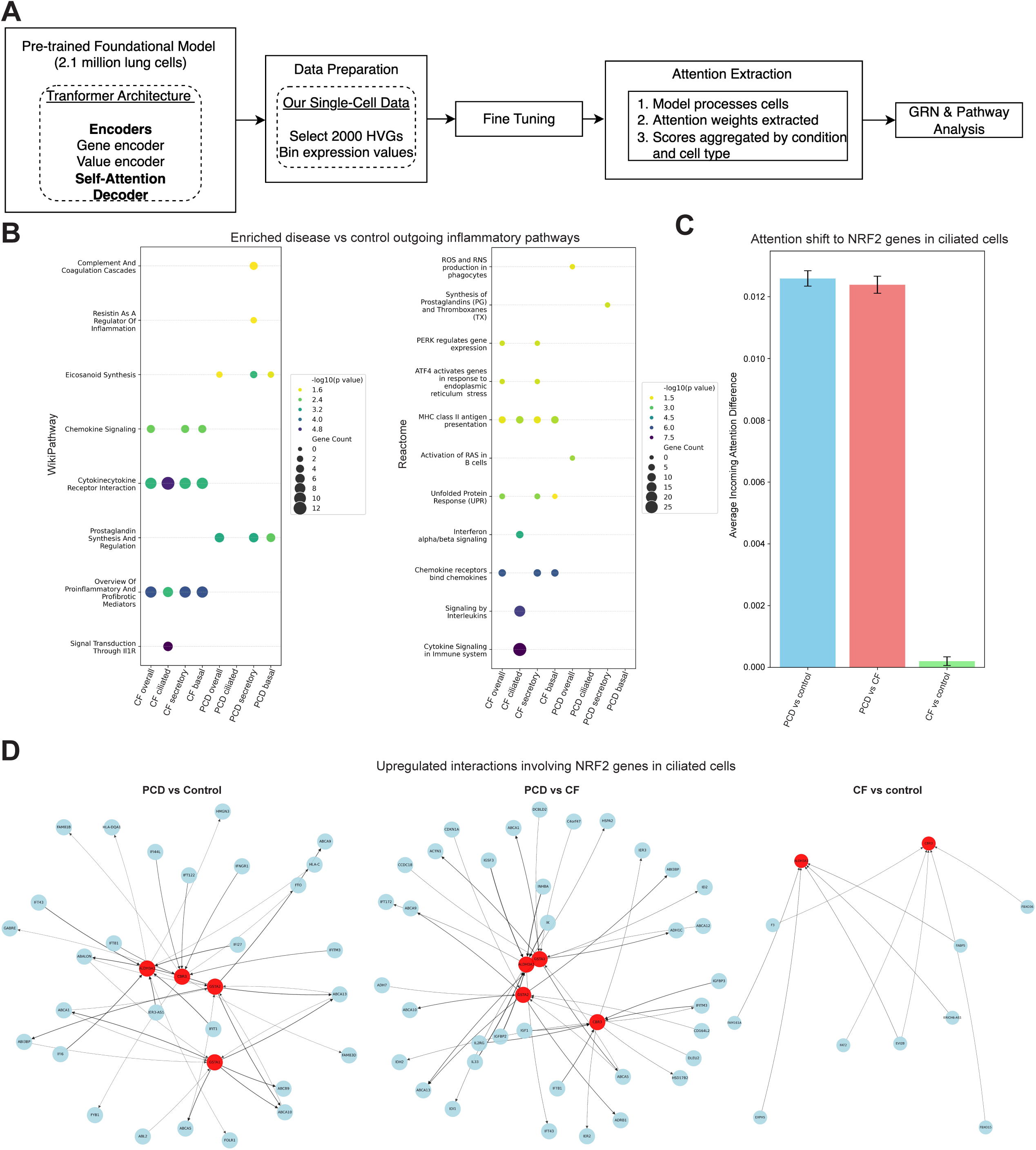
Gene regulatory network inference with a fine-tuned scGPT transformer model. **(A)** Schematic of the computational workflow for fine-tuning the scGPT lung foundation model and subsequent extraction and analysis of attention scores. **(B)** ORA of inflammatory pathways for the top 150 genes exhibiting the highest increase in ’outgoing attention’ in CF and PCD cells, each compared to controls. **(C)** Bar plot quantifying the differential ’incoming attention’ shift towards NRF2 pathway-associated genes in ciliated cells for three comparisons: PCD vs. Control, PCD vs. CF, and CF vs. Control. Error bar represents the standard error of the mean (SEM). **(D)** Gene regulatory networks (GRNs) depicting differential attention interactions (PCD vs. Control, PCD vs. CF, and CF vs. Control) involving NRF2-associated genes in ciliated cells. Edge weight is proportional to the differential attention score.

Using this approach, we performed overrepresentation pathway analysis on the top 150 genes with the greatest increase in “outgoing attention” vs control, which represents a gene’s regulatory influence over a network (**Figure 5B**). CF cells showed significant enrichment for pathways related to cytokine, interleukin, interferon signaling and unfolded protein response (UPR) activation. Further enrichment of chemokine signaling in secretory cells and immune response pathways indicated a heightened, pro-inflammatory state, aligning with our differential gene expression analysis. In contrast, the top enriched pathways in PCD were related to the arachidonic acid metabolism and the synthesis of pro-inflammatory eicosanoids, along with the complement pathway.

**Figure 5.**
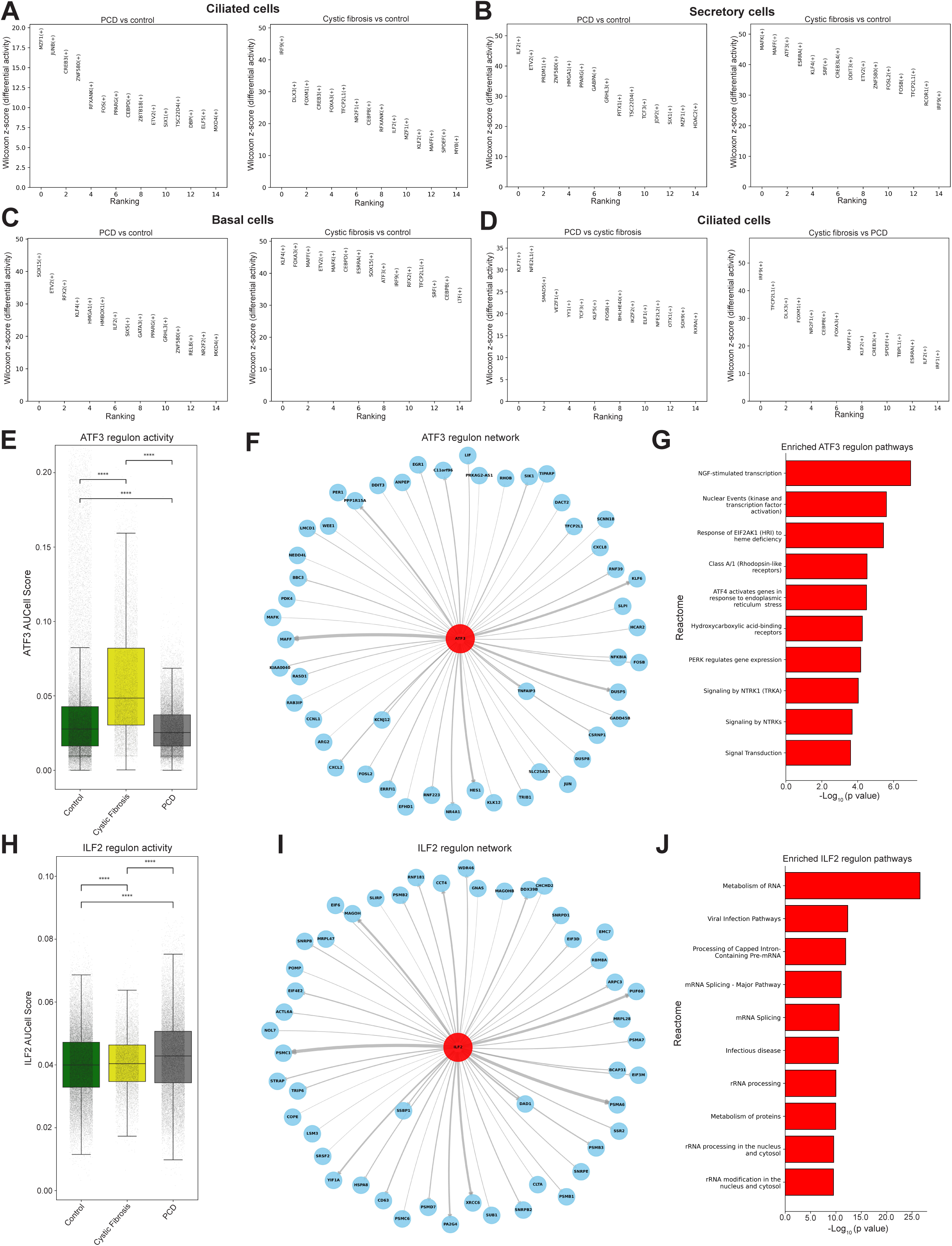
Gene regulatory network inference with SCENIC. **(A-D)** Summary plots display differentially active transcription factor regulons identified from AUCell scores. Regulons were compared using the Wilcoxon rank-sum test, filtered for an adjusted p-value (p < 0.05), and ranked by z-score. Plots show the top 15 regulons for: **(A)** ciliated cells, comparing PCD vs. Control (left) and CF vs. Control (right); **(B)** secretory cells, comparing PCD vs. Control (left) and CF vs. Control (right); **(C)** basal cells, comparing PCD vs. Control (left) and CF vs. Control (right); **(D)** a direct comparison between PCD and CF ciliated cells. **(E)** Boxplots quantifying the AUCell score for the ATF3 regulon across conditions. Statistical significance was determined by a pairwise Wilcoxon rank-sum test; significance levels: *p < 0.05, **p < 0.01, ***p < 0.001, ****p < 0.0001; ns, not significant. **(F)** Network visualization of the *ATF3* regulon, displaying the top 50 target genes ranked by grnboost2 correlation score, where edge weights are proportional to the score. **(G)** Top 10 enriched Reactome pathways from an overrepresentation analysis (ORA) of all target genes within the ATF3 regulon. **(H)** Boxplots quantifying the AUCell score for the ILF2 regulon across conditions, with statistical significance noted as in **(E)**. Network visualization of the ILF2 regulon, with edge weights proportional to the grnboost2 correlation score. **(J)** Top 10 enriched Reactome pathways from an ORA of all target genes within the ILF2 regulon.

Moreover, analysis showed an incoming attention shift towards NRF2 genes (*GSTA1, GSTA2, ALDH3A1* and *CBR1*) in ciliated PCD cells when compared to both CF and controls (**Figure 5C**). While only a slight increase in CF ciliated cells was observed.

To dissect the specific gene-gene interactions driving this increased signature, we generated the GRNs depicting the most significant connections involving these NRF2-asssociated genes in PCD and CF ciliated cells (**Figure 5D**). The NRF2 network in PCD versus control was defined by a strong crosstalk with the innate immune system. The NRF2 hubs receive elevated attention from interferon-stimulated genes (*IFIT1*, *IFI6*, *IFI44L*, *IFITM3* and *IFI27*) as well as the interferon gamma receptor *IFNGR1* and intraflagellar transport genes (IFT43, *IFT81* and *IFT122*) and in turn influenced antigen presentation machinery (*HLA-DQA1* and *HLA-C*). NRF2’s regulation of interferon signaling is well documented in respiratory disease (22). In contrast, when compared to CF, the PCD network was characterized by its integration with growth factor (*IGFBP2* and *IGFBP3*) and specific interleukin (*IL33* and *IL2RG*) signaling pathways (**Figure 5D**). Consistent gene-gene interactions with the NRF2 hubs and intraflagellar transport genes were once again observed (*IFT172*, *IFT43* and *IFT81*). These PCD networks showed a recuring pattern of interactions between *GSTA1* and *GSTA2* with ABC transporters, which may suggest increased activity of glutathione-mediated detoxification in PCD compared to controls and CF. Less NRF2 gene interactions were observed in the CF vs control network, with *GSTA2* and *GSTA1* lacking any increased interactions (**Figure 5D**).

#### Transcription factor programs that drive PCD and CF disease states

To identify key transcription factors (TFs) and their target gene networks (regulons) driving the disease states, we performed SCENIC regulatory network inference analysis (23). Regulon activity was quantified for each cell using an AUCell score (area under the curve of regulon genes ranked by expression within a cell). A higher AUCell score indicates that a larger portion of regulon genes are among the most highly expressed genes in that cell, reflecting stronger regulon activity. Differentially active TFs were then identified between the different conditions by comparing these scores (**Figure 5A-D**).

The transcriptional program in CF was defined by a strong stress and elevated immune signaling program. In CF secretory cells, the network was dominated by regulons of the endoplasmic reticulum (ER) stress and unfolded protein response (UPR) (**Figure 5B**). Evidenced by the enrichment of the *ATF3* regulon, a key transcription factor downstream of the PERK-eIF2α-ATF4 branch, that regulates the terminal UPR, apoptosis and autophagy (24). Deeper analysis of the global *ATF3* regulon confirms its role in connecting these cellular processes. The analysis revealed a high AUCell score for the *ATF3* regulon in CF compared to PCD and controls (**Figure 5E**). To understand the functional nature of this response, we analyzed the genes that comprise the *ATF3* regulon (**Figure 5F**). This network of target genes includes canonical genes of the terminal UPR (*DDIT3*), pro-apoptotic factor *BBC3* and pro-inflammatory chemokines (*CXCL8*, *CXCL2*). Furthermore, to confirm the collective biological function of this active regulon, a pathway enrichment analysis was performed on the entire set of *ATF3* target genes and revealed the PERK-ATF4 signaling axis in response to ER stress as a top pathway (**Figure 5G**), while a direct comparison between CF and PCD, revealed *ATF4* as a top scoring regulon (**Supplemental Figure 4B**). The pro-inflammatory state of CF secretory cells was substantiated by the enrichment of the *KLF4* and AP-1 complex (*FOSL2, FOSB*) regulons (**Figure 5B**). In CF ciliated and basal cells, the signature was characterized by a strong interferon response program, governed by the activity of interferon regulatory factor *IRF9* and was further marked by the ER stress response through *ATF3* and *CREB3* activation (**Figure 5A and 5C**).

The transcriptional program in PCD was similarly defined by inflammation and stress, but through a different set of active regulons. In secretory cells, the signature was marked by *ILF2*, a multifunctional protein that regulates interleukin-2 and is hypothesized to act as a stress support gene, enhancing DNA repair capacity, maintaining RNA translation homeostasis and suppressing apoptosis (25) (**Figure 5B and 5H**). Further analysis of the *ILF2* regulon showed its target genes were enriched for pathways related to infection, RNA metabolism and protein homeostasis (**Figure 5I and 5J**).

Analysis of multiciliated cells showed that the AP-1 complex (*FOS, JUNB*) was enriched in ciliated PCD cells rather than the interferon-centric program observed in ciliated CF cells (**Figure 5A**). A direct comparison between the disease states further confirmed the unique oxidative stress signature in PCD, identifying *NFE2L2* as a top differentially active regulon in multiciliated and secretory cells (**Figure 5D and Supplemental Figure 4A**). A deeper analysis into the NF-kB signaling pathway, an established source of exaggerated inflammation in CF (26), revealed that *NKB1* and *NFKB2* regulons were significantly increased in CF and downregulated in PCD cells compared to controls (**Supplemental Figure 4C and 4E**). Interestingly, antioxidant response regulators, *MAFF* and *MAFK* were differentially active in CF compared to PCD and controls in all cell types (**Figure 5A-D and Supplemental Figure 4B**).

#### Increased cellular responses to stress in CF and PCD

To evaluate and compare the cellular stress responses of CF and PCD to controls, we calculated the standardized mean difference (Cohen’s d) for a panel of stress-related pathway modules scores from the Reactome database (16). This approach quantifies the magnitude of the difference for each module in diseased cells relative to controls. The effect sizes, visualized in a heatmap (**Figure 6A**), revealed shared and divergent signatures. The markers driving the increased stress programs were then identified through differential gene expression analysis (**Figure 6B**)

**Figure 6.**
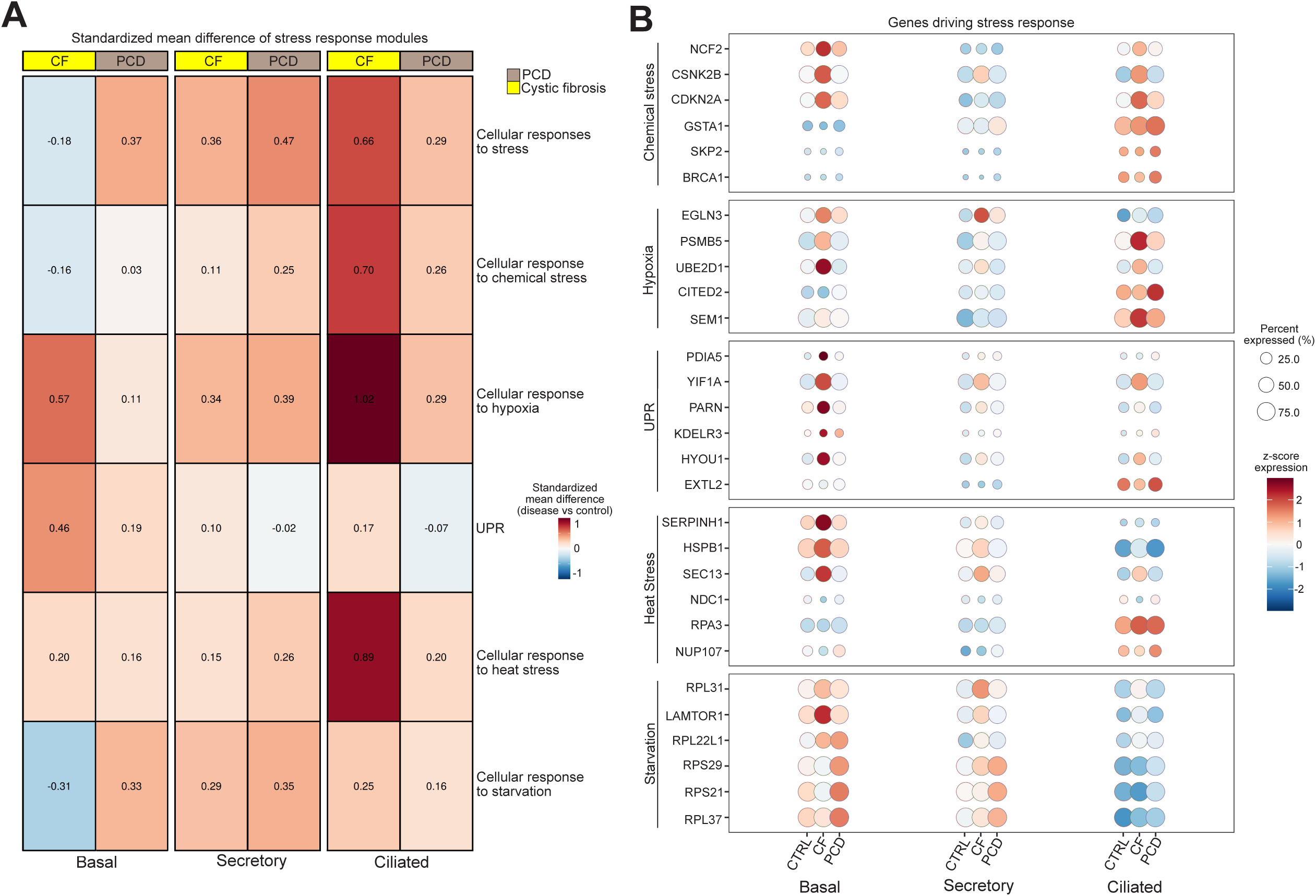
Cellular responses to stress. **(A)** Heatmap of standardized mean difference (Cohen’s d) for cellular stress module scores. This plot compares the effect size of pathway activity in diseased (CF and PCD) versus control cells across the major cell types. Rows represent the indicated stress-related pathways from Reactome. Columns are grouped by cell type and split by condition. Red indicates a positive Cohen’s d (higher pathway activity in disease), while blue indicates a negative Cohen’s d (lower activity in disease). The color scale is centered at zero, and the specific Cohen’s d value is overlaid on each cell. **(B)** Dot plot depicting the percent expression and z-score expression of the top 6 differentially expressed genes comparing CF and PCD against the pooled controls for the respective genes in each stress module. DEGs were ranked by Log_2_ fold change and filtered by adjusted p-value (p < 0.05). The plot is split by condition and visualized across the major cell types: basal, secretory and ciliated.

The overarching cellular responses to stress module was elevated in the secretory and ciliated cells of both PCD and CF, suggesting a broad stress phenotype (**Figure 6A**). The cellular response to chemical stress module, which manages the consequences of chemical insults and damage from reactive oxygen species, was similarly elevated in both conditions, though with a stronger effect size in CF ciliated cells compared to the slighter increases in PCD (**Figure 6A**). The CF signature was characterized by increased expression of *NCF2* in basal and ciliated cells, a component of the ROS-producing NADPH oxidase complex (27) and *CSNK2B*, a protein kinase involved in DNA damage repair and stress signaling (**Figure 6B**) (28). In PCD, the signature was defined by NRF2-associated detoxification enzyme *GSTA1* in ciliated cells (**Figure 6B**).

Analysis of more specific stress pathways revealed clear differing signatures for each disease. CF cells showed a pronounced upregulation of the cellular response to the hypoxia module (**Figure 6A**), particularly in ciliated cells, with increased expression of ubiquitin-proteosome components *PSMB5* and *UBE2D1* (**Figure 6B**). PCD showed a more modest increase in the hypoxia module, primarily restricted to ciliated and secretory cells, and was supported by the increased expression *CITED2*, a known negative regulator of *HIF1A* signaling (29).

The unfolded protein response (UPR) module score was increased in CF basal cells. This finding is consistent CF biology, where misfolded CFTR can induce ER stress (30) and was characterized by the upregulation of key ER-resident proteins, including chaperone *HYOU1*, protein foldase *PDIA5* and ER-retrieval receptor *KDELR3* (**Figure 6B**). Cellular response to heat stress was also exhibited in CF ciliated cells (**Figure 6B**), while the cellular response to starvation module, which manages adaptations to nutrient deprivation, displayed an elevated signal in PCD basal and secretory cells, supported by the expression of ribosomal genes (**Figure 6B**). These data demonstrate the stress profiles are distinct, with CF characterized by hypoxia and UPR pathways, while PCD showed more modest levels for the cellular stress signatures.

## Discussion

Cystic fibrosis (CF) and Primary Ciliary Dyskinesia (PCD) both result in chronic sinopulmonary disease from impaired mucociliary clearance, and often PCD care is derived from experience with cystic fibrosis. The distinct transcriptional signatures of both conditions at a single-cell resolution have not yet been directly compared (1). Here, we asked if the different genetic origins of CF (channelopathy) and PCD (ciliopathy) lead to divergent or convergent cellular stress and inflammatory responses in the airway epithelium. Using single-cell RNA sequencing, we compared the transcriptomes of cultured primary airway epithelial cells against each other and against healthy individuals. Our analysis revealed that while both diseases share a clear elevation in immune signaling that implicates neutrophil recruiting cytokines, they may be driven by fundamentally different transcriptional programs.

The CF signature was characterized by heightened interferon and chemokine signaling, along with evident unfolded protein response activity. Conversely, the PCD signature was defined by the notable activation of the NRF2 pathway and a more modest inflammatory profile involving neutrophil chemotaxis.

Consistent with prior observations (7), we found that differentially expressed genes in multiciliated PCD cells compared with control cells were associated with downstream effectors of glutathione and the NRF2 pathways, now with a larger and more diverse set of PCD variants. Here, we also observe a similar increase in this subset of oxidative protection genes and pathways in PCD when compared to CF. The NRF2 pathway is a critical cellular defense mechanism where, under conditions of oxidative stress, it activates an array of antioxidant and cytoprotective genes (31). As we previously reported, NRF2 is particularly relevant to PCD and is required to maintain normal cilia length and beat frequency by protecting the ciliary axoneme from oxidative damage (7).

We have hypothesized, that in PCD variants with residual dynein arm structures, like *DNAH5*, the remaining motors exhibit increased activity in a compensatory effort, resulting in an elevated oxidative burden from increased ATP metabolism and subsequent NRF2 activation (7). Our results now demonstrate this as a more conserved transcriptional response, with PCD samples with variants in *CCDC39* and *CCDC40* exhibiting upregulation of NRF2 effectors, including *GSTA2*. The activation of *GSTA2*, a key phase II detoxification enzyme that conjugates electrophilic compounds with glutathione, signifies the engagement of an adaptive cytoprotective program in response to a heightened state of oxidative stress in PCD multiciliated cells (32). Using single cell GPT (scGPT) gene regulatory network analysis, we further show the centrality of this pathway by revealing its increased attention score and suggest it acts as a key hub integrating antioxidant defenses with innate immune and growth factor signaling.

The loss of *CFTR* function in CF is known to inherently create a pro-oxidative environment due to a fundamental redox imbalance (33) and is exacerbated by the defective transport glutathione to the air-liquid surface (34, 35). Compounding this intrinsic vulnerability, it has been demonstrated that this state of chronic oxidative stress paradoxically leads to a suppression of NRF2 and *GSTA2* activity at both the protein and transcriptional level (36, 37). Our findings align with this response, with the NRF2 signature and *GSTA2* activity decreased in CF compared to PCD. The adaptive NRF2-GSTA2 activation in PCD compared to the inherently disrupted antioxidant system in CF represents a key molecular distinction in their respective pathophysiology, which may contribute to distinct mechanisms of cellular damage.

Our comparative analysis confirms that CF and PCD are defined by elevated inflammatory profiles, consistent with established reports of neutrophil-dominated airway inflammation in both conditions (38). Although both disorders share features like chronic respiratory infections and excessive neutrophil influx, the underlying lung pathology in PCD is far less characterized than in CF. We found that much of this inflammatory activity was localized to secretory cells, in which both conditions showed strong upregulation of pro-inflammatory cytokines, including the potent neutrophil chemoattractant *CXCL8*. However, further analysis revealed that the transcriptional programs driving this inflammation are different. The cytokine and neutrophil signaling signatures were more pronounced in CF, a finding corroborated by our scGPT analysis, and the aggregated expression score of a cytokine signaling module. A more profound distinction was the identification of a uniquely heightened interferon response in CF cells, supported by the activity of regulatory factor *IRF9*.

In contrast, we identified an increased transcriptional signature for arachidonic acid metabolism in PCD, which may provide a potential mechanistic link between ciliary dysfunction and the characteristic neutrophilic inflammation. Leukotriene B4, a potent neutrophil chemoattractant (39) and prostaglandins, which modulate neutrophil activation and resolution (40), both demonstrated increased transcriptional signaling in CF and PCD. The divergence in inflammatory drivers was also evident at the level of transcription factor activation; while the pro-inflammatory AP-1 complex was active in both conditions, key downstream regulons such as *NFKB1* and *NFKB2* were upregulated in CF but not in PCD. The inability of misfolded CFTR to reach the cell surface prompts intrinsically high levels of NF-kB, which mediates the release of pro-inflammatory cytokines (26).

Another central distinction we observed in the cellular stress programs of CF and PCD is the activation of the unfolded protein response (UPR) in CF, which appears to be a strong driver of its pathophysiology. While the UPR is composed of three main signaling branches, our data points to the PERK-eIF2α-ATF4 signaling axis as a transcriptionally active component of the response (30). The SCENIC analysis, which is a computational tool for simultaneous gene regulatory network reconstruction (23), reveals high activity of *ATF4* and *ATF3* regulons. The activation of the PERK branch of ER stress can signal through NF-kB (41, 42), which in turn drives the expression of a suite of cytokines, including *CXCL8*, which was associated in *ATF3*’s regulon, contributing to chronic inflammation in CF.

Transcriptional results suggested, however, that despite the antioxidant deficiencies in CF, ciliated cells still exhibited a pronounced internal compensatory response program, potentially involving *NCF2*-mediated ROS signaling and the stress-response kinase *CSNK2B*. In contrast, *GSTA1* was a key driver specific to PCD ciliated cells, consistent with a coordinated, NRF2-driven adaptive program. The suppression of NRF2 in CF may be tied to the high activity of *MAFF* and *MAFK,* whose regulons were differentially activated across all CF major cell types. While small Maf proteins are required partners for NRF2, activation, their overabundance can promote the formation of repressive homodimers that block NRF2-mediated transcription (43).

The cellular response to hypoxia was similarly increased in both diseases with responses strongly augmented in CF ciliated cells, along with a modest increase in PCD ciliated cells. The thick stagnant mucus produced in CF, due to ASL dysregulation is known to create hypoxic environments (44). The less pervasive response in PCD, was characterized by the *HIF1A* negative-regulator, *CITED2*. This suggests that while PCD cells may experience some level of oxygen deprivation, it may not be the driving stressor that it is in CF, and the cellular responses appear to be actively restrained. The adaptive stress response profile in PCD is potentially demonstrated by the activation of *ILF2*, a multifunctional protein hypothesized to act as a stress support gene (25, 45, 46).

ScRNA-seq analysis is a powerful tool for investigating cell-type specific gene expression. However, standard differential gene expression (DEG) testing carries a risk of false positivity due to pseudoreplication bias (47). Our primary findings were supported by convergent computational evidence from our fine-tuned scGPT model. This approach is not susceptible to the same statistical biases as DEG testing. Identifying the upregulated NRF2 pathway in PCD and the neutrophil dominant inflammatory state of CF cells through scGPT further validate the DEG analysis results.

Our study integrated single-cell transcriptomic data from multiple cohorts, employing a tiered analytical approach to maximize the statistical power while controlling for potential batch effects. Direct comparisons between disease states utilized the fully integrated dataset, while disease versus control analyses were restrained within their respective cohorts; local PCD versus local controls and public CF versus public controls, to minimize site-specific technical variation. For computationally intensive network analyses, we used the larger, robust public CF cohort. We directly compared our two local CF samples to the five public CF samples. This analysis revealed differences in cellular proportions and the number of DEGs between the two cohorts, reinforcing the presence of inter-cohort variability and the importance of controlling for culture condition and cell preparations between different sites.

In summary, our comparative single-cell analysis of primary airway cells from patients with CF and PCD provides insight into their shared and divergent programs. While both diseases exhibit a clear inflammatory state, our findings indicate they may be driven by fundamentally distinct pathophysiology. We show that the CF signature is defined by a notable elevation in chemokine signaling, UPR-centric stress response and heightened interferon activity. Whereas the PCD signature is characterized by the activation of an NRF2-driven antioxidant response and unique metabolic profile.

## Methods

### Airway epithelia cell culture

Human nasal epithelial cells were obtained from PCD and CF patients or healthy participants after obtaining confirmed consent approved by the Institutional Review Board at Washington University. Deidentified nasal epithelial cells were retrieved using a cytology brush biopsy of the inferior nasal turbinate. Primary airway cells were processed, expanded in culture, and then differentiated using ALI conditions in Transwell (Corning Inc.) as previously described (7, 9).

### scRNA-seq

ScRNA-seq data from PCD subjects with variants in *CCDC39* and *CCDC40*, as well as variants in *DNAH5* were previously published and made available (GSE275070 and GSE272189, respectively) (7, 9). Overall, samples were locally acquired from 7 healthy individuals, 7 PCD patients (variants: 4 *DNAH5*, 2 *CCDC39*, 1 *CCDC40*) and 2 CF patients (2 f508del). In short, primary nasal epithelial cells were harvested on ALI days 28-30 after cilia presence was confirmed by direct visualization and immunofluorescent staining using a cilia marker. Library preparation and sequencing were conducted at the Genome Technology Access Center at Washington University. At least 10,000 cells per sample were processed using a Chromium Controller (10X Genomics) for single-cell capture, with 7,000 to 10,000 cells sequenced per sample. cDNA was prepared post-GEM generation and barcoding, followed by GEM-RT reaction and bead cleanup, adhering to manufacturer protocols. Purified cDNA underwent 11-13 amplification cycles before SPRI select bead cleanup. Samples were run on an Agilent Bioanalyzer to measure cDNA concentration. Gene expression libraries were prepared using 10X Genomics Chromium Single Cell 3’ Reagent Kits (v3.1 Chemistry Dual Index), adjusting PCR cycles based on cDNA concentration.

The Chromium Next GEM Single Cell 3’ Kit v3.1, Chromium Next GEM Chip G Single Cell Kit, and Dual Index Kit TT Set A were used for sample preparation. Library concentrations were accurately determined using qPCR with the KAPA Library Quantification Kit (KAPA Biosystems/Roche) to achieve appropriate cluster counts for the Illumina NovaSeq6000 instrument. Libraries were normalized and sequenced on a NovaSeq6000 S4 Flow Cell with a 50×10×16×150 sequencing recipe per the manufacturer’s protocol, targeting a median sequencing depth of 50,000 reads per cell. Paired-end sequencing reads were processed by Cell Ranger (10X Genomics software, version 8.0.0), with reads aligned to GRCh38 (version 90) for genome annotation, demultiplexing, barcode filtering, and gene quantification. Barcodes with less than 10% of the 99th percentile of total unique molecular identifiers (UMIs) per barcode were excluded from analysis. Gene barcode matrices were generated by counting UMIs for each gene (rows) in individual cells (columns).

Additional CF single-cell sequencing data were retrieved from two separate published datasets comprising of airway cultured cells. FASTQ files for five samples were extracted from the Gene Expression Omnibus (GEO) series GSE150674 (8). Three ALI controls and two F508del variants made up the samples. Another five ALI samples were extracted from GEO Series GSE191061, comprising of two controls and three CF samples (10). The FASTQ files were then processed using the Cell Ranger count pipeline to generate gene-cell UMI count matrices.

### scRNA analysis

Seurat R package version 5.0 was used to process and analyze the single-cell RNA sequencing datasets, which included global unsupervised clustering to identify distinct cell populations and evaluate compositional differences (48). Quality control filtering was performed to remove cells with low gene counts (minimum 200 genes detected), high mitochondrial gene content (generally exceeding 20-25%), or high ribosomal protein gene content (generally exceeding 20-35%) were excluded. Genes detected in less than three cells were also removed from the analysis. Following QC, potential doublets were identified and removed from each sample individually using the Doublet Finder R package with a presumed doublet rate of 7.5% (v2.0) (49). Separate integrated datasets were created for the local samples/controls and the public CF samples/controls. A combined analysis, integrating all samples and controls was also generated to allow for a direct comparison between PCD and CF. To correct for batch effects across samples, while preserving biological heterogeneity, data normalization was performed using the multiBatchNorm function from the batchelor R package version 3.21 (50). Principal component analysis (PCA) was run individually on each sample layer. Datasets were then integrated using canonical correlation analysis (CCA). Dimensionality reduction using uniform manifold approximation and projection (UMAP) was performed on the first 30 dimensions of the integrated CCA reduction for visualization. Cell clustering was achieved using Seurat’s FindNeighbors function followed by the Leiden algorithm (leidenAlg v1.15) for identifying groups of interconnected cells (51). Initial cell type annotation was performed by examining the expression of canonical marker genes. Clusters exhibiting mixed marker expression or requiring finer distinction underwent iterative subclustering and were selectively annotated using known markers from previous transcriptomic analysis (7).

#### Differential Expression Testing

To identify differentially expressed genes (DEGs) between conditions within the major cell types, the FindMarkers function within Seurat was used. The Wilcoxon Rank Sum test was used to compare expression levels between specified groups. All comparisons were performed using data filtered by DEG identification using a log-fold change threshold of 0.5. For defining significant DEGs for downstream analyses, a Bonferroni-corrected adjusted p-value < 0.05 was used, combined with a detection in at least 10% of cells in one of the comparison groups. Unless directly comparing PCD and CF, differential expression analysis was confined to testing between diseased cells and their batch-specific controls.

#### Over-Representation Pathway Analysis

Pathway over-representation analysis (ORA) was conducted using the clusterProfiler R package (v3.2.1) to investigate the biological functions associated with differentially expressed genes (52). For each comparison of interest, the DEG lists were verified using the HGNChelper R package (v0.8.15) and mapped to ENTREZ Gene IDs (53). ORA was performed on the resulting list. Enrichment against the WikiPathways 2024 and the Reactome database was carried out (16, 20, 54).

#### Single cell GPT analysis

To infer and compare gene regulatory networks (GRNs), the pre-trained scGPT lung model (2.1 million cells) was fined-tuned using our integrated dataset, which consisted of the top 2000 highly variable genes with expression values binned into 51 categories (21). The fine-tuning process used an 80/20 train/validation split, a 15% masking probability, mean squared error (MSE) loss, an Adam optimizer (1 × 10^−5^ learning rate) and early stopping based on validation loss to achieve the best performing model. The model was fine-tuned for 19 epochs and achieved a validation loss of 0.1078, closely tracking the training loss of 0.1080. Attention weights, representing gene-gene interaction strengths, were extracted from the final transformer layer of the fine-tuned model. These cell-specific attention scores underwent rank normalization and were averaged across attention heads before being aggregated by condition and cell type to generate average attention matrices. Differential attention matrices were then computed by subtracting these averaged matrices between conditions. Pathway ORA was performed on the top 150 genes with the most increased outgoing attention for each group. Significant pathways related to inflammatory and immune processes were then visualized. A specific analysis on NRF2-related genes in ciliated cells involved visualizing increased attention interactions via network plots and calculating the average incoming attention differences towards these genes.

#### SCENIC analysis

To identify active transcription factor-centered gene regulatory networks (regulons), we used the pySCENIC (v0.12.1) pipeline (23). Gene regulatory networks were inferred from the raw count matrix of the combined integrated dataset using GRNBoost2. The resulting co-expression modules underwent motif enrichment analysis using prune2df, which utilized motif ranking databases and annotations to identify direct genes with enriched motifs for each transcription factor. The activity of each regulon was then quantified using AUCell, (which measures regulon activity based on the expression ranking of their target genes) generating an AUCell matrix, which was added to the single-cell object. To identify differentially active regulons between conditions, the Wilcoxon rank-sum test with Benjamini-Hochberg correction was applied to the AUCell scores within major cell types for comparisons between conditions. Regulons were filtered by an adjusted p-value < 0.05 and ranked by the Wilcoxon z-score.

#### CellChat analysis

To investigate intercellular communication networks, with a focus on immune and inflammatory signaling, we used the CellChat R package (v2.2.0) (17). We created separate CellChat objects for Control, CF and PCD conditions from the normalized count matrix of the integrated dataset. The human ligand-receptor interaction database (CellChatDB.human) was used as the reference. ComputeCommunProb was utilized to infer the communication probabilities for each dataset with the “trimean” option. Pathway-level communication was then calculated. The three objects were then merged to facilitate a comparative analysis of signaling pathways across conditions. We focused on the upregulated immune signaling pathways in CF and PCD and used the rankNet function to compare their relative information flow. To further investigate the key signaling pathways within arachidonic acid metabolism, specifically prostaglandin, circle plots were generated using the netVisual_aggregate function. The ligand-receptor pairs contributing to the expanded prostaglandin signaling in PCD were then displayed using the netVisual_chord_cell function.

#### Gene Module Scores

To quantify the activity of key biological pathways, module scores were created using Seurat’s AddModuleScore function. This score aggregates the expression of a pre-defined set of functionally related genes on a per-cell basis. We generated scores for the NRF2 cytoprotective, Type I interferon, and Type II interferon signaling pathways using gene sets from WikiPathways (20). We also generated scores for the following pathways from the Reactome database, to comparatively analyze signaling in PCD and CF: Cytokine signaling in immune system, Cellular responses to stress, Cellular response to chemical stress, Cellular response to hypoxia, Unfolded protein response, Cellular response to heat stress and Cellular response to starvation (16). To quantify the magnitude of difference in the stress-related pathways’ activity between conditions, the standardized mean difference (Cohen’s d) was calculated for each module score. Within each major cell type, the effect size was computed by comparing the disease states against the pooled control group, using the respective means, standard deviations and cell counts. The resulting matrix was visualized as a heatmap. To then visualize the genes driving these pathways, the top three differentially expressed genes between each disease state and controls were selected, ranked by Log_2_ fold change and plotted.

## Tables

**Table 1.** Characteristics of local donors. Table shows the clinical characteristics of all local donors involved in the study.

