## Supplementary figures and images for "Comparative Single-Cell Transcriptomics Uncovers Shared and Distinct Molecular Signatures in Cystic Fibrosis and Primary Ciliary Dyskinesia"

### Supplemental Figure 1

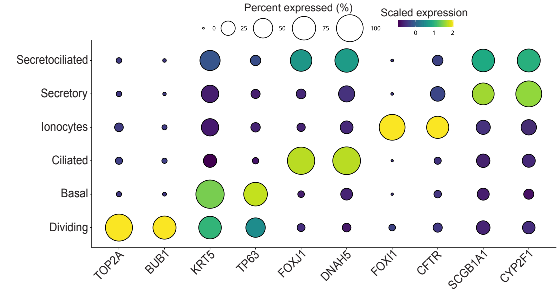

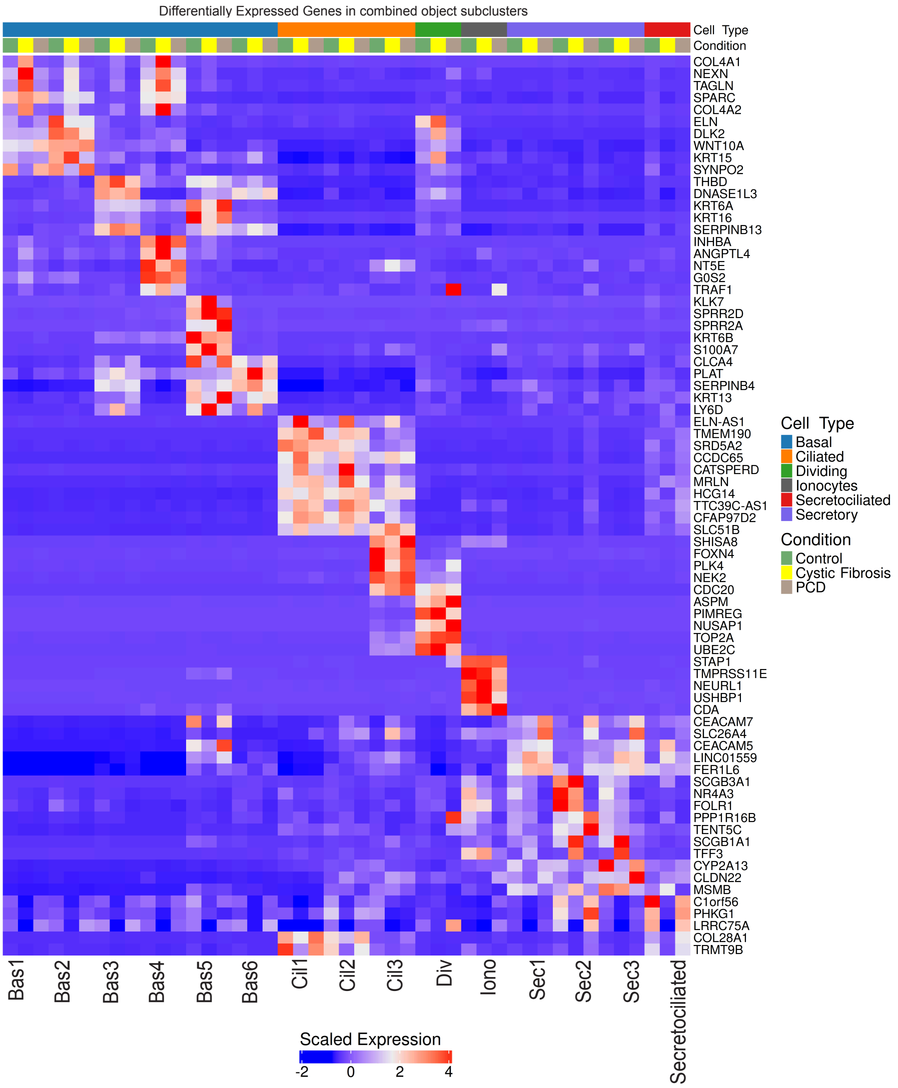

### Supplemental Figure 2

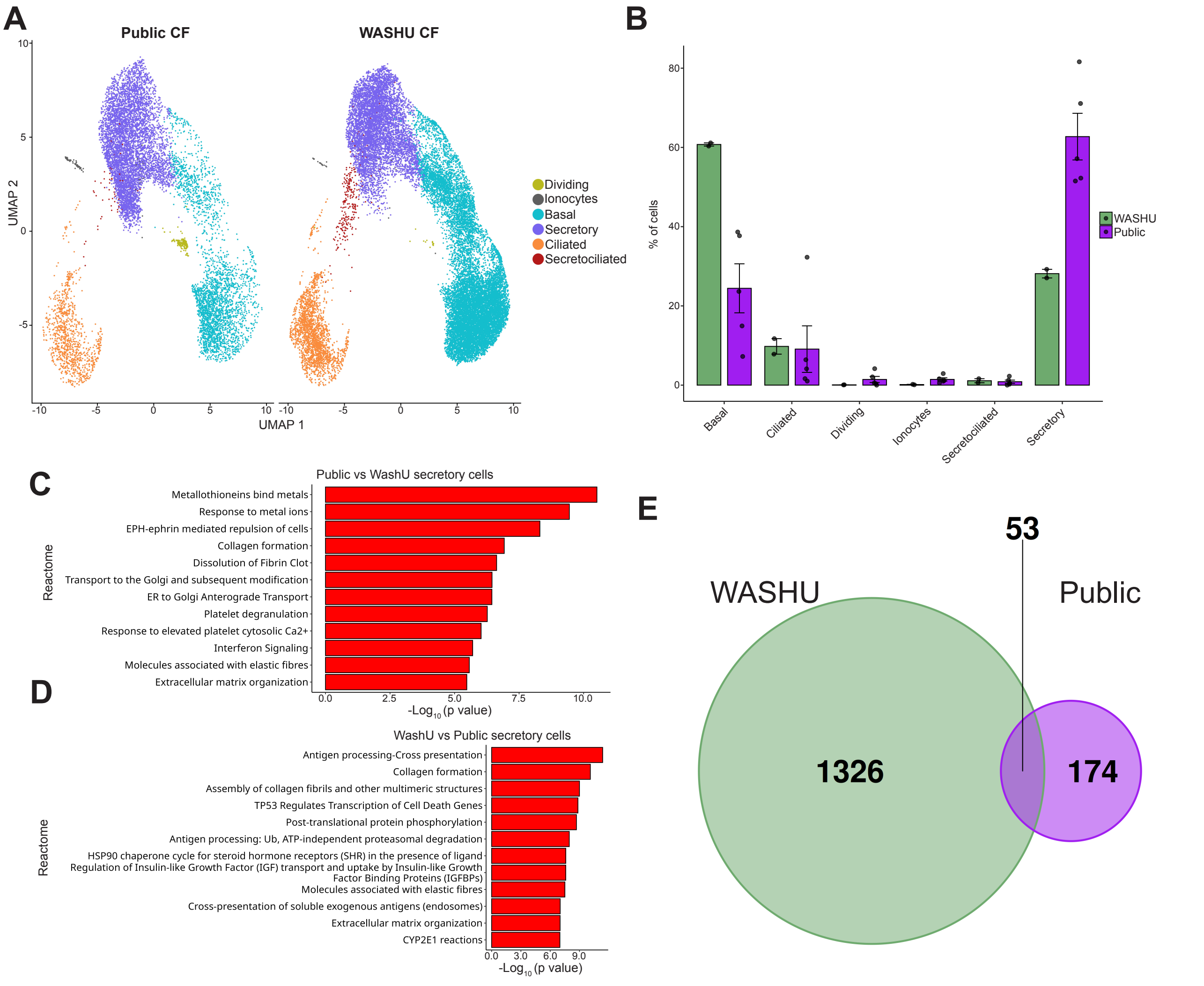

### Supplemental Figure 3

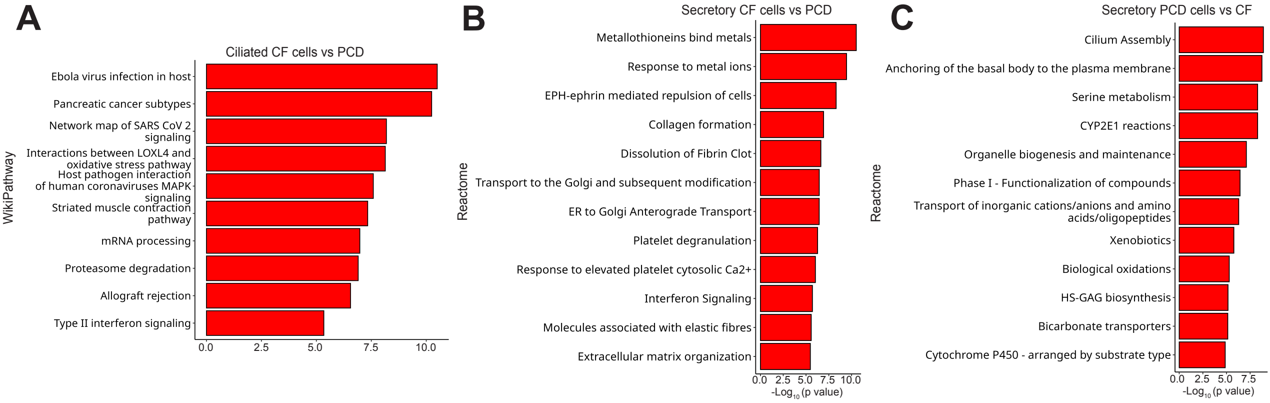

### Supplemental Figure 4

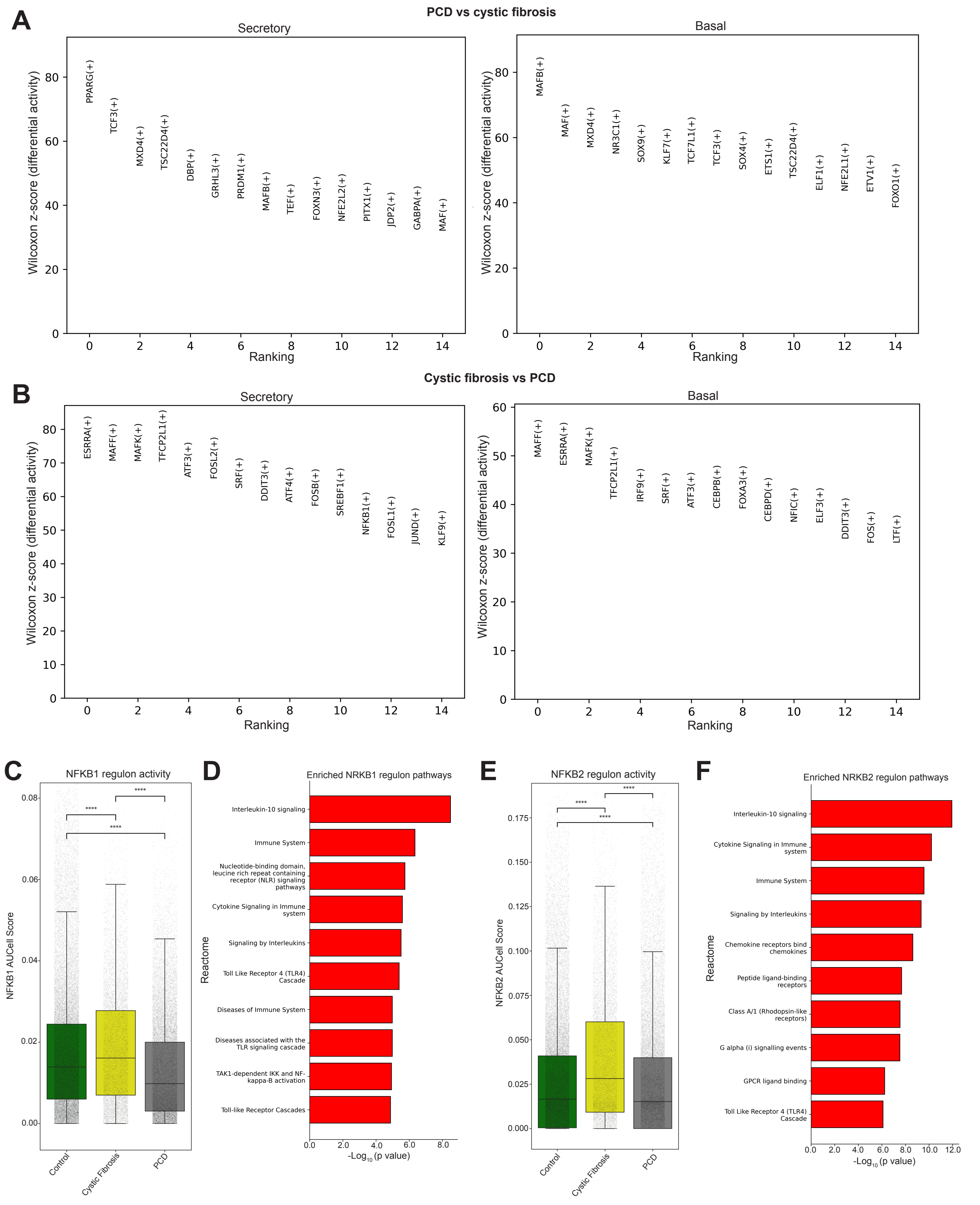
